# Multiphoton tomographic fluorescence lifetime imaging microscopy – TomoFLIM

**DOI:** 10.64898/2026.09.11.750847

**Authors:** Liam Collard, Ani Augustine Jose, Fahmida Rahman, Rogelio Garcia Aguirre, Conor Treacy, Tommy Pallett, Sian Culley, Simon Ameer-Beg, Simon Poland

## Abstract

Fluorescence lifetime imaging microscopy provides quantitative, concentration-independent contrast for probing molecular interactions, biochemical environments and cellular physiology. However, the requirement to acquire sufficient time-resolved photon statistics makes FLIM inherently slow, limiting its application to dynamic biological processes. In live-cell applications, including calcium signalling and vesicular trafficking, acquisition times can exceed the timescale of the underlying biology, causing temporal averaging, motion artefacts and loss of transient information. Methods that increase FLIM acquisition speed while preserving quantitative lifetime accuracy are therefore required.

We report on a high-speed, compressive, multiphoton fluorescence lifetime imaging technique (TomoFLIM). Two-photon fluorescence is excited using a line focus projected tomographically across the sample, while time-tagged fluorescence is acquired using time-correlated single-photon counting. Time-resolved fluorescence data are reconstructed using Lucy-Richardson deconvolution followed by a computationally efficient centre-of-mass method lifetime estimator. In addition, TomoFLIM Net, a physics-informed neural-network, directly reconstructs fluorescence intensity and lifetime from compressed time-resolved tomographic data. TomoFLIM was benchmarked against raster-scanned fluorescence lifetime measurements using calibrated fluorescence lifetime beads and biological specimens. We demonstrate imaging at compression ratios exceeding 90%, with Pearson correlation coefficients above 80% relative to reference images. A raster-scanned FLIM dataset acquired in 60 s was reproduced using TomoFLIM in 3.75 s, representing a 16-fold increase in frame rate and equivalent reduction in accumulated dark counts. TomoFLIM Net recovered distinct experimental bead lifetime populations, demonstrating a direct route from compressed measurements to quantitative lifetime maps. TomoFLIM therefore offers significant potential for rapid live-cell imaging of dynamic biological processes, including deep within turbid biological specimens.

## Introduction

Fluorescence lifetime imaging microscopy (FLIM) provides quantitative information on cellular and molecular processes ^1–4^, but the acquisition of high-resolution lifetime images remains inherently slow, imposing a trade-off between spatial information and imaging speed that limits the observation of dynamic biological processes. By measuring the arrival times of the fluorescence photons relative to a pulsed excitation source, the decay rate of the excited fluorophore can be calculated. One of the most widely adopted implementations of FLIM combines raster-scanning microscopy with reverse start-stop time-correlated single-photon counting (RSS-TCSPC). Here, the arrival time of each detected photon is recorded relative to the subsequent excitation pulse, enabling fluorescence decay curves to be reconstructed with picosecond temporal resolution^5^. RSS-TCSPC requires sufficient photon statistics to be collected at each pixel. Under the low excitation powers required to minimise photobleaching, photodamage, and pulse pile-up, particularly during multiphoton excitation, this often necessitates pixel dwell times ranging from tens to hundreds of microseconds. This is mitigated by either measuring lifetime in the frequency domain^6^ or by using time-gated detection using ICCDs^7^, CMOS^8^, and SPAD arrays^9^ to acquire wide-field lifetime images. However, these techniques do not achieve the excellent temporal resolution of RSS-TCSPC, which is considered the gold standard for lifetime measurement. Consequently, the acquisition of high-resolution raster-scanned lifetime images may require tens of seconds^10^, preventing the observation of many dynamic biological processes and limiting throughput in live-cell imaging.

The development of SPAD arrays^11^ incorporating time-to-digital converters for TCSPC has enabled fluorescence lifetime measurements to be parallelised both spatially and spectrally, substantially increasing imaging throughput. SPAD line sensors have been used for time-resolved single-photon detection across multiple spectral channels^12^, to capture spatially resolved signals along one axis to achieve confocal line scanning^13^ and, through light-field optical encoding, to recover three-dimensional lifetime information^14^. In addition to line detectors, the combination of 2-dimensional array detectors and beam shaping optics such as spatial light modulators and diffractive optical elements has been used to achieve microscopes parallelised along both the x and y spatial axes for confocal^15,16^, wide-field^17^ and multiphoton^18,19^ FLIM. These approaches can provide substantial improvements in FLIM frame rates, however depending on detector architecture, limitations can include wavelength-dependent instrument response functions, photon detection efficiency, and the requirement to preserve spatial correspondence between the excitation pattern and individual detector elements. This spatial correspondence becomes increasingly difficult to maintain in strongly scattering samples, where fluorescence emitted at depth can be redistributed before reaching the detector.

More recently, compressive imaging approaches utilising single-pixel detectors^20–23^ have gained traction as a means of increasing FLIM acquisition speed. These techniques encode spatial information into a reduced set of measurements from which an image can be computationally reconstructed. In Ma et al.^24^, a DMD applied pseudorandom masks to the collected fluorescence. The fluorescence was projected onto a streak camera, enabling readout of the lifetime for each pattern. In Ghezzi et al.^25^, the authors used a DMD to apply masking on the excitation light to encode randomness. This system has since been extended to support computational 3D multispectral FLIM, demonstrating the potential of compressive approaches for volumetric and multidimensional imaging^26^. In parallel, a hyperspectral time-resolved single-pixel FLIM platform using Hadamard structured illumination in both excitation and collection arms, enabling quantitative wide-field fluorescence lifetime imaging and *in vivo* FRET measurements over a large field of view has been demonstrated^27^.The impact of different sampling bases for this system was also subsequently investigated^28^. More recently, the same group introduced Net-FLICS, replacing the computationally intensive compressed sensing inversion and pixel-wise lifetime fitting with a deep neural network capable of directly reconstructing fluorescence intensity and lifetime images from raw single-pixel measurements^29^. In addition, projection-based approaches have also enabled reconstruction of three-dimensional images based on data projected onto a gated CCD ^30^. Although these studies demonstrate the potential of compressive imaging for FLIM, existing single-pixel compressive FLIM implementations have predominantly focused on wide-field excitation. In parallel, compressive approaches have been used to accelerate intensity-based multiphoton microscopy. Liquid-crystal spatial light modulators and DMDs have been used to generate three-dimensional beamlet patterns^31,32^, which are then used as a basis for compressive multiphoton imaging. By projecting a modulated line beam across the sample^33^ kilohertz-frame-rate two-photon imaging has also been demonstrated. These approaches demonstrate that structured multiphoton excitation can substantially reduce the number of measurements required to recover spatial information.

In this work, we introduce a hybrid approach, tomographic fluorescence lifetime imaging microscopy (TomoFLIM), which combines single-pixel TCSPC detection with patterned multiphoton illumination and computational reconstruction. This approach retains the key advantages of raster-scanned multiphoton microscopy, including optical sectioning, efficient excitation and non-descanned fluorescence detection, while reducing the number of spatial measurements required for lifetime-image reconstruction. A line focus is projected across the sample at a series of angular orientations, generating time-resolved tomographic projections that retain a direct physical relationship to the underlying spatial distribution. This provides a structured alternative to pseudorandom compressive measurements and removes the requirement for a spatially resolved detector such as a SPAD array. In addition to conventional computational reconstruction, we investigate whether a physics-informed neural network approach, TomoFLIM Net, can directly reconstruct fluorescence intensity and lifetime from compressed time-resolved TomoFLIM projections. This provides an alternative, extremely fast computational route for interpreting TomoFLIM data without requiring independent reconstruction of multiple temporal moments. The system is shown to be capable of imaging biological samples at compression ratios of over 90% with a Pearson’s correlation coefficient (PCC) of over 0.8 relative to raster-scanned reference measurements while retaining the principal fluorescence lifetime distributions.

## Results

### Outline and Principles of TomoFLIM

TomoFLIM combines the benefits of random compressive FLIM, ordered raster-scanned imaging, and parallelised FLIM with single-pixel detectors and TCSPC as illustrated in **Figure 1a**. A multiphoton laser and a cylindrical lens, mounted in a high-speed rotation stage, generate a line focus which is projected orthogonally to its long axis across the sample using galvo mirrors. For a raster-scanned lifetime image containing *N* × *N* × *t_bins_* data points an equivalent TomoFLIM projection contains *N* × *N_θ_* × *t_bins_* data points where *N_θ_* is the number of angles in the projection (which is typically one to two orders of magnitude lower than *N*) and *t_bins_* is the number of temporal bins. A circular aperture is positioned at the focal point of the cylindrical lens effectively flattening the excitation profile. In addition, it ensures that the field of view along each projection axis is the same.

**Figure 1.**
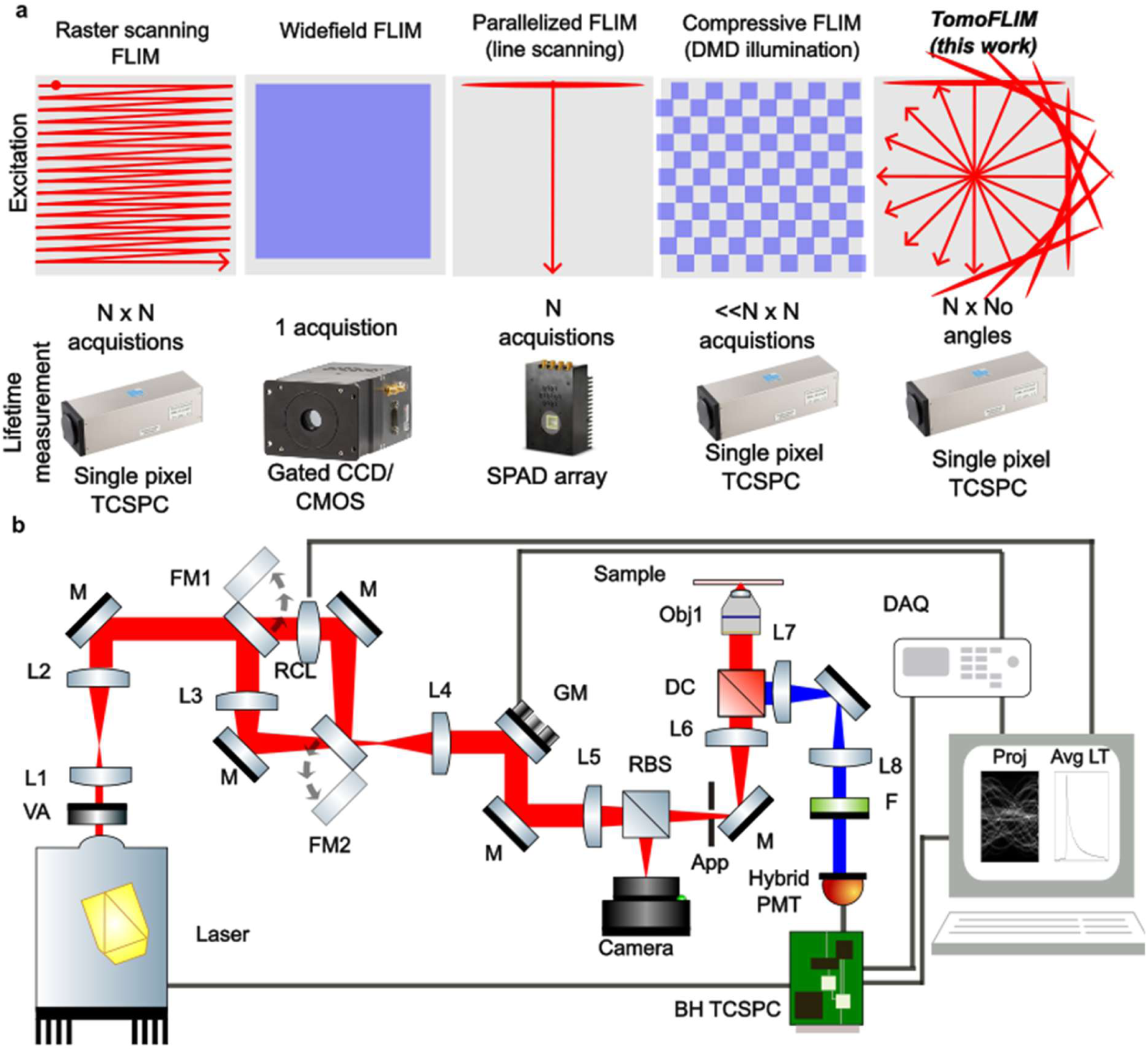
– **a** Summary of fluorescence lifetime imaging techniques and TomoFLIM. **b** Microscope design and hardware L – lens, M-mirror, FM – flip mirror, RCL – rotary cylindrical lens, GM - dual axis galvomirrors, RBS – removable beam splitter, App – aperture, DC – long-pass dichroic, Obj – microscope objective, F – filter, TCSPC – Becker and Hickl SPC -830, DAQ, data acquisition device. The computer illustrates the live readout of tomographic projection and average transient.

The arrival times of the collected fluorescence photons are measured using a single hybrid PMT detector TCSPC system. For each line focus position, a fluorescence decay is acquired, from which a fluorescence lifetime image may be reconstructed. The developed system also includes a conjugated raster-scanned imaging arm to enable comparison of TomoFLIM with raster-scanned RSS-TCSPC FLIM. Between raster-scanned imaging and TomoFLIM, the overall light dose exposure was equalised to ensure comparable datasets, by increasing the power by a factor of 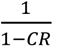 where *CR* is the compression ratio. The count rates were carefully monitored so as to ensure that we avoid the pulse pile-up regime. This results in approximately the same number of fluorescent counts in each image; however, because of the shorter measurement time, TomoFLIM records fewer dark counts. A complete description of the TomoFLIM system utilised in this work and shown in **Figure 1b** is described in the Materials and Methods section.

### Interpreting TomoFLIM data

The TomoFLIM projection produces a tensor indexed by an angular index *i_θ_*, spatial index *i_z_* and arrival time *t*. The average transient (i.e. the summed data across the spatial dimensions) and the total tomographic projection (data summed across the temporal dimension) are displayed live in MATLAB and illustrated in **Figure 1b**. Reconstructing lifetime images from TomoFLIM projections is a complex mathematical and computational problem involving both image data decompression and decay modelling. **Figure 2** illustrates two approaches we have applied to interpret TomoFLIM data. Initially, data were histogrammed from macro-time photon events into 2^B^ temporal channels (typically 256 where B = 8) and we take a time-point by time-point and subsequent curve-fitting approach, as illustrated in **Figure 2a-d**. To reconstruct a fluorescence lifetime image from the individual time-channel tomographic projection, a Lucy-Richardson reconstruction (see methods) was applied on the tomographic projection using the previously published method^33^, for each time channel. The reconstructed image was imported into TRI2 software^34^, and a mono-exponential decay curve was fitted (Levenberg–Marquardt algorithm) to recover the lifetime and intensity and shown in **Figure 2d**. This data was imported back into MATLAB for visualisation. The same fitting algorithm was applied to the raster-scanned images to establish the ground truth. Whilst curve fitting can provide the gold standard estimation of lifetime measurement for raster-scanned images, we identified several issues when attempting to extend this to TomoFLIM data. Firstly, for the earlier temporal bins, the reconstruction algorithm is capable of faithfully reconstructing the resultant image. However, at later temporal bins, far fewer photons are available from this to reconstruct an image. This results in greater uncertainty of the Lucy-Richardson reconstruction at these bins and weakens confidence in fluorescence lifetime estimation. To mitigate this, we attempted to down-sample our data from 256 to 32 bins. Whilst this resulted in some improvement, ultimately, we found that this resulted in broad lifetime histograms compared to ground truth raster-scanned imaging. Furthermore, the down sampling worsens the temporal resolution. In addition to the scientific issues, the computation is time consuming and required 32 Lucy-Richardson computations.

**Figure 2.**
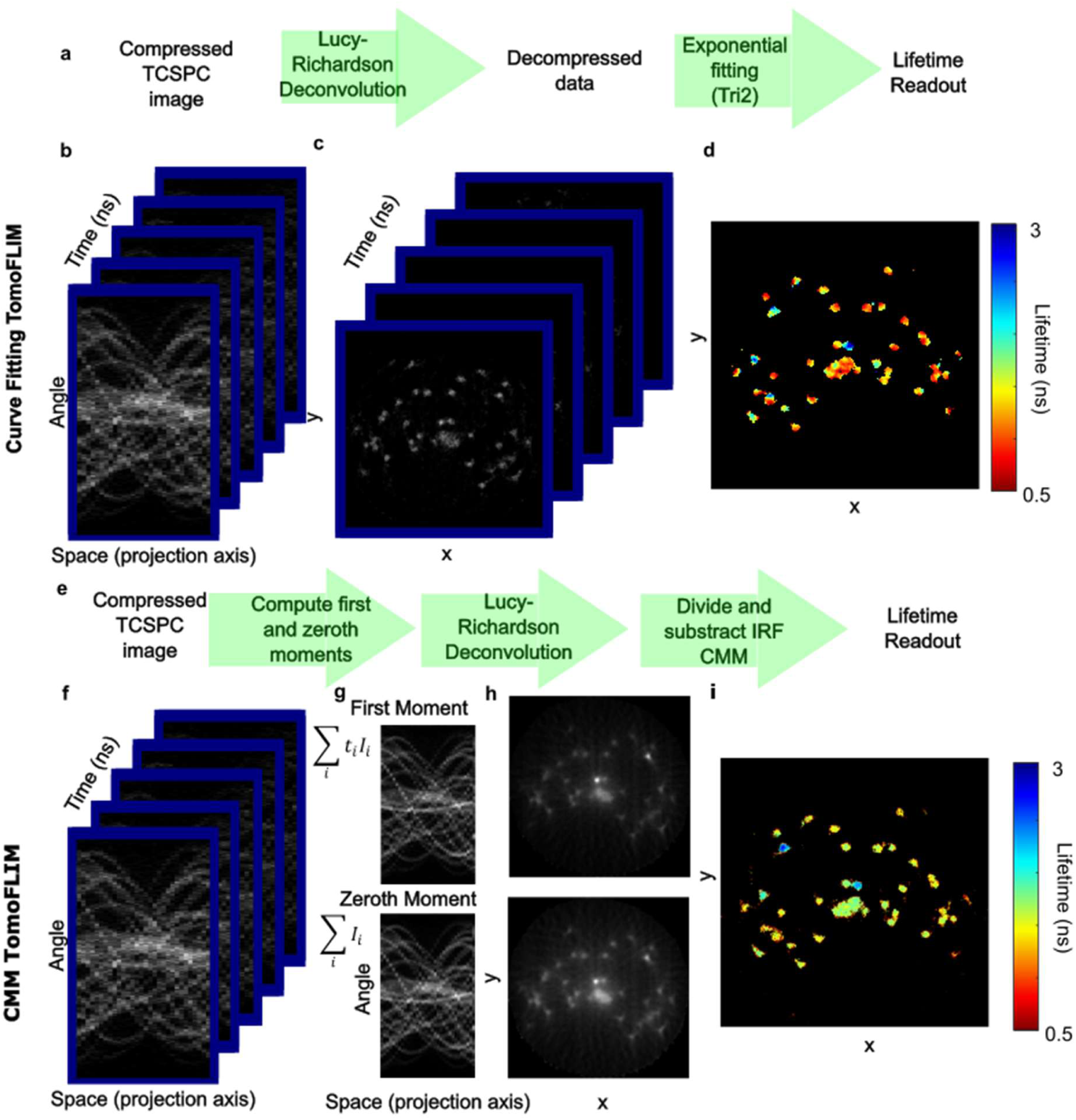
– **a** Illustration of the curve fitting processing pipeline from tomographic, TCSPC projections to reconstructed compressive lifetime images. **b** Tensor of tomographic projections across the temporal axis. **c** Lucy–Richardson reconstructions of each tensor. **d** A resultant lifetime image generated by fitting a single exponential to the data in panel c. **e** Illustration of the centre of mass method for reconstruction of lifetime from TomoFLIM data. **f** Tensor of tomographic projections across the temporal axis. **g** The zeroth and first moments of the TomoFLIM data. **h** Lucy–Richardson reconstructions of the zeroth and first moments. **i** Resultant lifetime image.

To alleviate these issues, we have developed a centre of mass method (CMM) for compressive lifetime reconstruction (**Figure 2e-i)**. Recall that for a lifetime decay curve 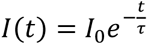 where τ is the fluorescence lifetime and *T* is the measurement window, the CMM is given:

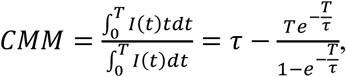

Which approximates to τ for *T* » τ. Here, we approximate 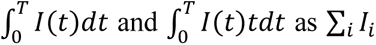 and ∑*_i_t_i_I_i_* which are calculated on the raw TomoFLIM data, where *i* indexes the temporal bins and *I_i_* is the measured intensity at each bin. The Lucy-Richardson algorithm is applied to both quantities independently, and they are divided to yield the CMM. Finally, the CMM of the instrument response function (IRF) is subtracted to yield the true lifetime^35^. This exhibits several advantages over the curve fitting approach. Firstly, the signal to noise is high and of the same order in each Lucy-Richardson computation. In addition, only two Lucy-Richardson computations are required and therefore within our implementation can provide readout within a few minutes. Approximation of the data to a single exponential decay is an acceptable compromise in the majority of sensor applications.

To evaluate the performance of these techniques, raster-scanned images and TomoFLIM projections were acquired from fluorescence lifetime calibration beads and Etzold-stained *Convallaria majalis* stem (**Figure 3**). The TomoFLIM datasets were analysed using both curve fitting and the CMM approach and the resultant fluorescence lifetime images were compared with raster-scanned data analysed using conventional curve fitting. Although direct quantitative comparison is complicated by differences between the reconstruction and analysis pipelines, the benefit of the CMM approach is apparent. For fluorescence lifetime calibration beads (**Fig. 3a–c**), there is clear spatial correspondence between the TomoFLIM CMM and raster-scanned curve fit data. On the contrary, in the curve-fit data, the lifetime appears slightly shorter. Remarkably, the TomoFLIM-CMM produces two clearly defined populations within the histogram and outperforms the curve fitting applied on the raster-scanned image. We attribute this to a lower level of overall background and therefore increased SNR in TomoFLIM. The system was shown to produce approximately 250 dark counts per second. For a raster-scanned image with 60 seconds acquisition time, this produces ∼15000 dark counts. By contrast, an equivalent TomoFLIM projection taken with 32 angles would result in ∼1000. This trend is also exhibited in the images of *Convallaria majalis* shown in **Figure 3d–f**. We thus proceeded to analyse all relevant data with curve fitting applied to raster-scanned images and the CMM approach to TomoFLIM data.

**Figure 3.**
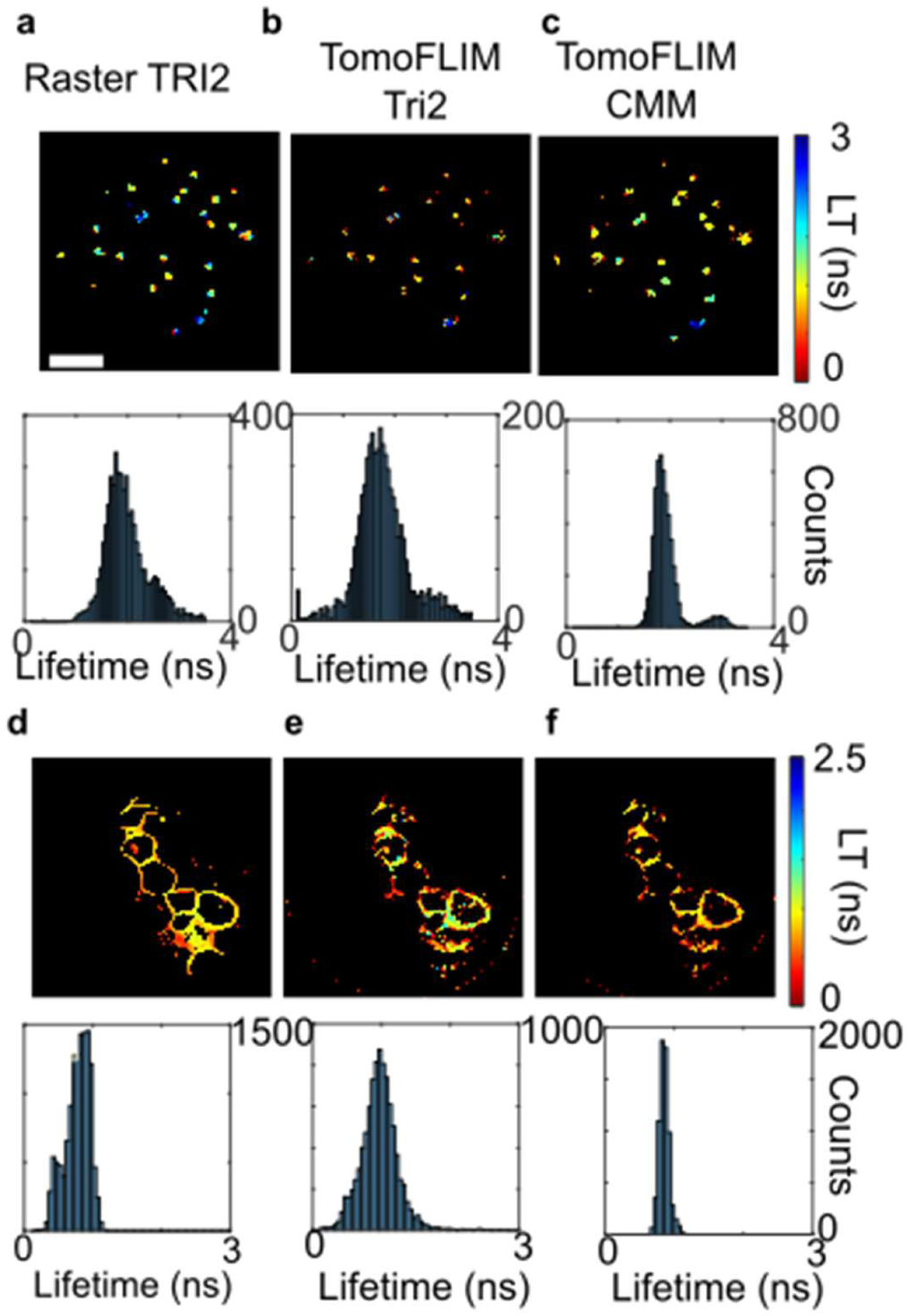
– Analysis of different techniques for interpreting TomoFLIM data. **a** Lifetime images of lifetime calibration beads taken with raster-scanned imaging and analysed with curve fitting. **b** Reconstructed lifetime images from TomoFLIM data and curve fitting. **c** Reconstructed lifetime images from TomoFLIM data and CMM. **d-f** Equivalent data for Convallaria.

### Evaluating compression ratio and image quality in TomoFLIM

To achieve an accurate image reconstruction, TomoFLIM, and other compressive imaging techniques require sufficient measurements relative to the sparsity of the image. In TomoFLIM, this can be achieved by increasing the number of projection angles in the tomographic projection, *N_θ_*. Our design, involving a cylindrical lens mounted in a high-speed rotation mount, enables real-time reconfiguration of this quantity. This can improve image quality but reduces the compression ratio and increases both measurement and reconstruction time. The simulations shown in the supplement (**Figures S1-S4**) show how image sparsity impacts reconstruction quality for intensity images for different *N_θ_* and compression ratio values. This is characterised in **Figure 4** for fluorescence lifetime with experimental data recorded on our TomoFLIM system. A raster-scanned image was recorded, and the system was switched to tomographic mode, and projections recorded with *N_θ_* = 32. This projection was subsampled to recover projections *N_θ_* = 16, 8, 4, 2. The resultant intensity and lifetime reconstructions are shown in **Figure 4**. The images were co-registered to correct small misalignments between tomographic and raster-scanned imaging modes, and the fidelity was quantified by measuring the Pearson’s correlation coefficient (PCC) between the raster-scanned image and reconstruction, which is inset on the images. PCC was chosen because it is less sensitive to image sparsity than pixel-wise error metrics and directly quantifies the agreement in spatial intensity (or lifetime) distributions between the raster-scanned and reconstructed TomoFLIM images^36^. For both intensity and lifetime measurement, *N_θ_* = 2 produces very poor reconstructions with very little similarity to the ground truth image. When *N_θ_* = 4 and *N_θ_* = 8, though some similarity is observed spatially (albeit with distortion), the lifetime values bear little resemblance to the ground truth image. With *N_θ_* = 16 and *N_θ_* = 32, both fluorescence intensity and lifetime reconstruction bear a strong correspondence to the raster-scanned image, and fluorescence lifetimes are spatially well separated. This can also be demonstrated by considering the histograms in **Figure 4**; as the compression ratio is decreased, the histograms become progressively narrower and more clearly defined. At *N_θ_* = 32, two well-defined populations are clear. This trend is also clearly reflected in the PCC of both the intensity and lifetime images.

**Figure 4.**
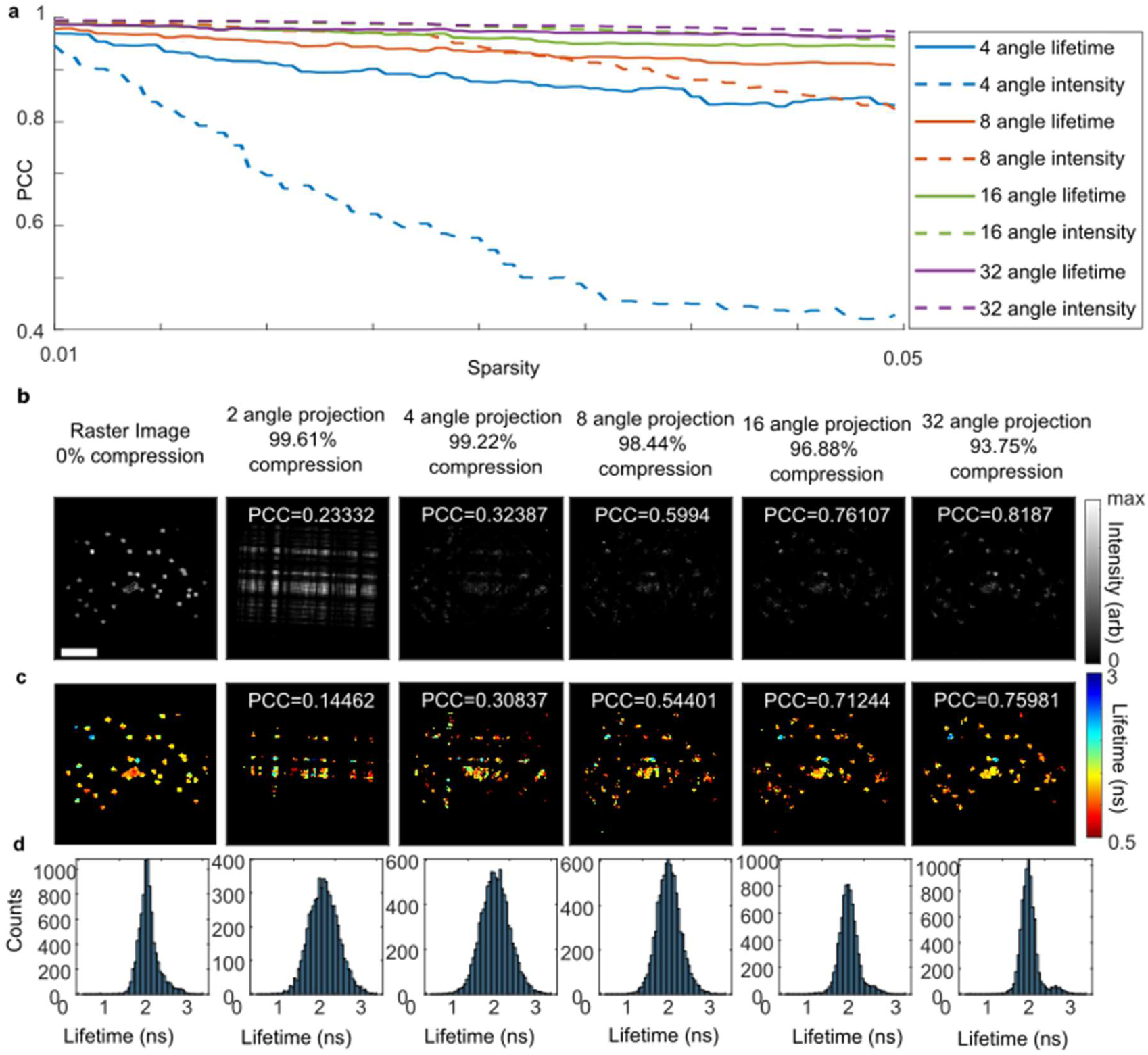
– Demonstration of tomographic reconstruction of intensity and lifetime images of lifetime calibration beads at different compression ratios. **a** Simulations showing the dependency of reconstruction quality of lifetime and intensity on compression ratio and image sparsity. **b** Intensity images for raster-scanned ground truth and reconstructions from experimentally measured TomoFLIM projections for N_θ_ = 2,4,8,16,32 (left to right). **C** Lifetime images for raster-scanned ground truth and reconstructions from experimentally measured TomoFLIM projections with N_θ_ = 2,4,8,16,32 (left to right). d Histograms of lifetimes of images shown in panel c.

To further examine the correspondence of compressive TomoFLIM acquisition to raster-scanned acquisition without compression, we imaged fluorescence lifetime calibration beads with both methods (**Figure 5)**. Across 4 unique fields of comparison, the PCC of the intensity images varied between 0.48 and 0.75. Most of the error is due either to small intensity differences of the beads in the image or slight alignment error due to imperfections in the image co-registration. Interestingly, the reconstruction tends to favour images with objects in the centre. These errors may correspond to the unevenness of the profile of the line beam used to excite the sample or to more complex field deformations which are often observed in high-numerical aperture lenses (likely small differences in apodization). Although a Gaussian line profile has been incorporated into the Lucy-Richardson algorithm, the true profile is more complex and varies with rotation due to small unavoidable aberrations and distortions within the system. Nevertheless, it is evident from the lifetime reconstructions, associated histograms and the PCC readout that the lifetime is faithfully reconstructed. Field aberration correction could be achieved with ground truth acquisition and affine transform correction but in this instance we chose not to do this and consider it a second-order effect.

**Figure 5.**
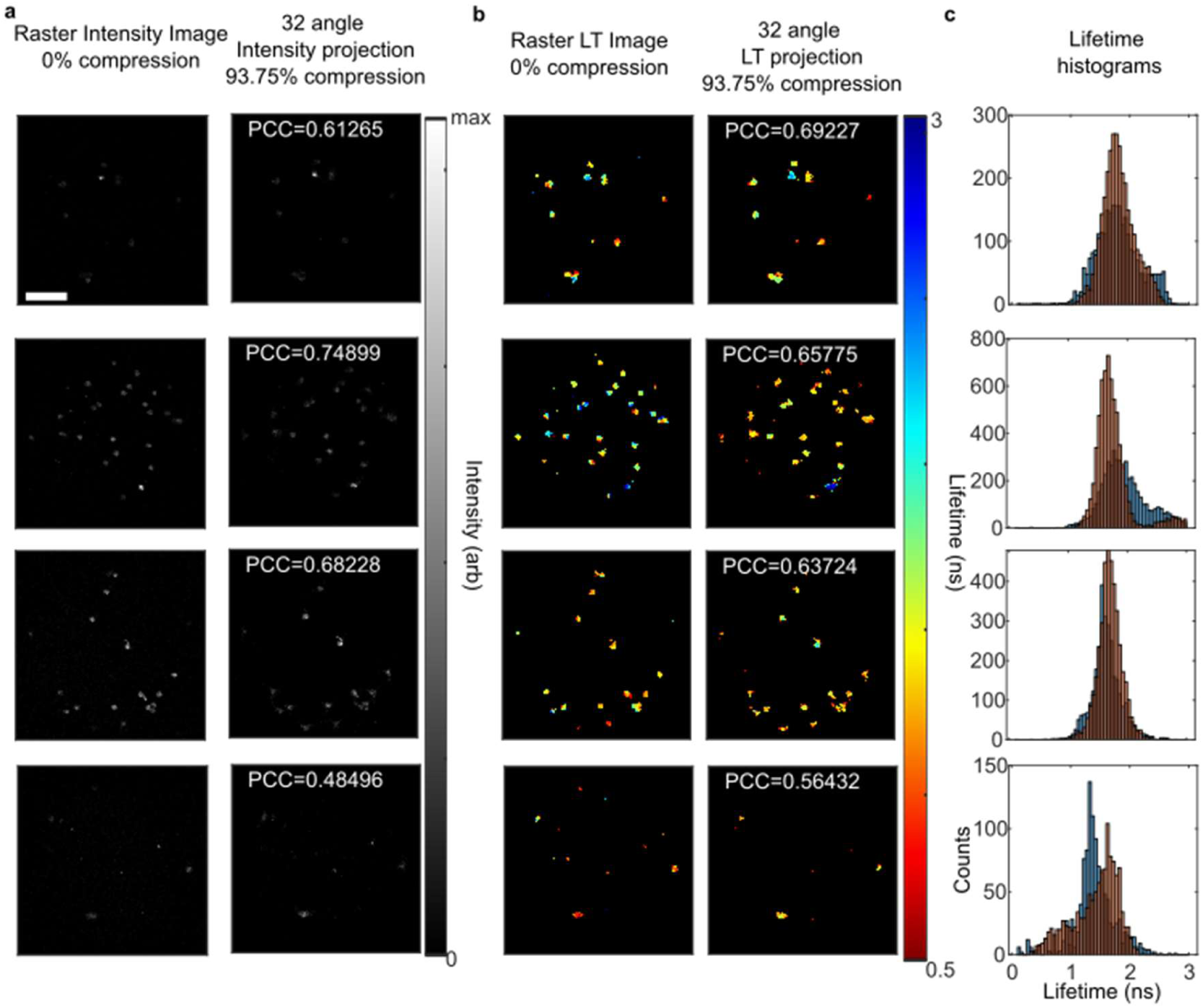
– Examples of tomographic images of lifetime calibration beads. **a** Intensity images left – raster-scanned ground truth, right tomographic reconstruction (32 angle projection) **b** Lifetime images left – raster-scanned ground truth, right tomographic reconstruction (32 angle projection). **c** Associated histograms for raster-scanned imaging lifetime (blue) and TomoFLIM (orange).

In order to examine the performance of TomoFLIM for more challenging biological samples, we acquired data for *Convallaria majalis* (**Figure 6)**. *Convallaria majalis* exhibits strong spatial variation in sparsity and is thus well suited to evaluate TomoFLIM, where the central core of the stem is substantially denser than the outlying cells. Clearly, the reconstruction depends strongly on sparsity; the fifth image (core of *convallaria*) has very low sparsity, and the PCC of both lifetime and intensity reconstruction is low (<0.6). For fluorescence intensity, the other images all exhibit PCC>0.7, suggesting a relatively high degree of similarity. The lifetime PCC exceeds 0.6 in five of the six fields and clearly captures spatial trends in fluorescence lifetime variation across the Convallaria. The histograms remain well aligned, and the TomoFLIM data produces extremely sharp fluorescence lifetime histograms due to the exceptional signal-to-noise ratio.

**Figure 6.**
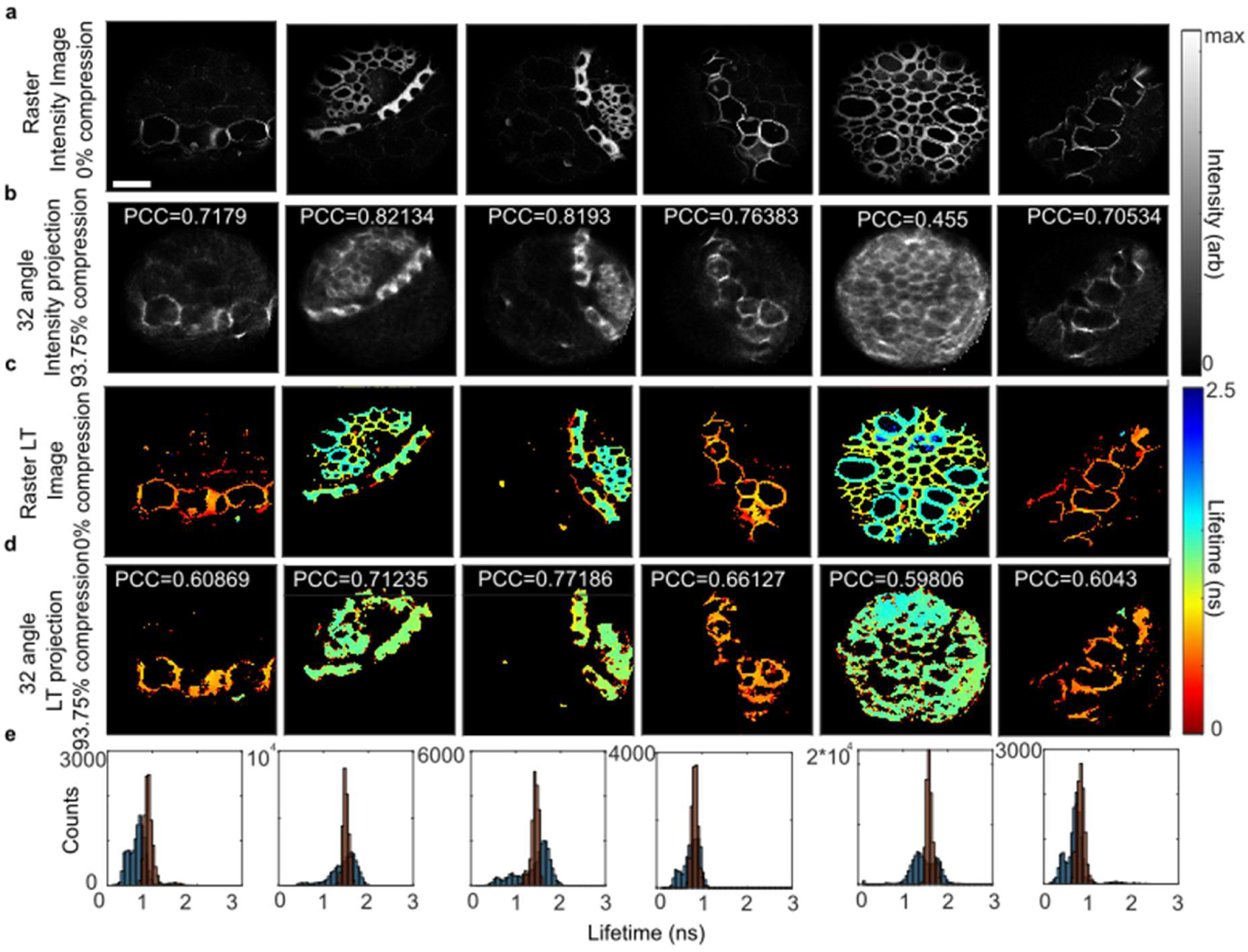
– Comparison of raster-scanned imaging and TomoFLIM for imaging Convallaria majalis. **a** Intensity images left – raster-scanned imaging. **b** TomoFLIM reconstruction of intensity (32 angle projection) **c** Lifetime images from raster-scanned imaging data measured with curve fitting. **d** Corresponding TomoFLIM lifetime reconstructions using CMM. **e** Associated histograms for raster-scanned imaging lifetime (blue) and TomoFLIM (orange).

### Learning to unscramble compressed TomoFLIM data using a physics-informed neural network

Although centre-of-mass reconstruction provided robust lifetime estimation from TomoFLIM data, it requires independent reconstruction of the zeroth and first temporal moments, which can be computationally time consuming. We therefore investigated whether a physics informed neural-network-based reconstruction could directly recover fluorescence intensity and lifetime from time-resolved tomographic data. The resulting network, *TomoFLIM Net*, was trained using simulated (512 by 512 by 256) images generated from a single representative raster-scanned TCSPC image recorded on our reference arm. Each simulated object was assigned an independently sampled mono-exponential lifetime, and the corresponding TCSPC decay was convolved with the experimentally measured IRF before projection through the 32-angle illumination patterns.

Rather than supplying the full time-resolved projection cube directly to the network, the data were represented by ten physically interpretable image-space features. These comprised an integrated intensity image, a normalised first temporal moment image, and eight temporally gated measurements, each obtained by the backprojection of the corresponding tomographic projection. This representation retained both spatial and temporal information while substantially reducing the dimensionality of the CNN input. The final network used an encoder-decoder architecture with channel widths of 16, 32, 48 and 64, learned stride-2 downsampling, two dilated bottleneck convolutions, transposed-convolution upsampling and additive skip connections. Layer normalisation, Leaky-ReLU activation and bottleneck dropout were incorporated before simultaneous relative-intensity, fluorophore-probability and continuous-lifetime outputs were generated (**Figure 7a**). Training and validation losses decreased consistently over 250 epochs, indicating stable optimisation without evidence of divergence (**Figure 7b**). On simulated data, TomoFLIM Net recovered both the spatial distribution of fluorescent objects and their continuous lifetime contrast. On the fixed 125-scene test data, the network achieved a lifetime mean absolute error of 0.0540 ns, lifetime root-mean-square error of 0.0990 ns, lifetime Pearson correlation coefficient of 0.9943, and intensity Pearson correlation coefficient of 0.9036 (**Figure S6**). Representative internal-benchmark images showed strong correspondence between the ground-truth and CNN-reconstructed intensity and fluorescence lifetime distributions (**Figure S6a**). Across the complete test data, predicted fluorescence lifetimes closely followed the ground-truth values over the simulated 0.5–4.0 ns lifetime range, with the pixel-wise measurements distributed closely around the line of unity (**Figure S6b**). The lifetime mean absolute error remained low over the investigated range of fluorescent-pixel sparsities, although a modest increase in error was observed as the image became less sparse (**Figure S6c**). These results demonstrate that TomoFLIM Net can recover continuous fluorescence lifetime values not explicitly represented as discrete training classes and remains robust over a broad range of image sparsities. Importantly, the lifetime output was accompanied by a fluorophore-probability map, allowing lifetime values to be restricted to pixels in which the network predicted sufficient fluorescent signal.

**Figure 7.**
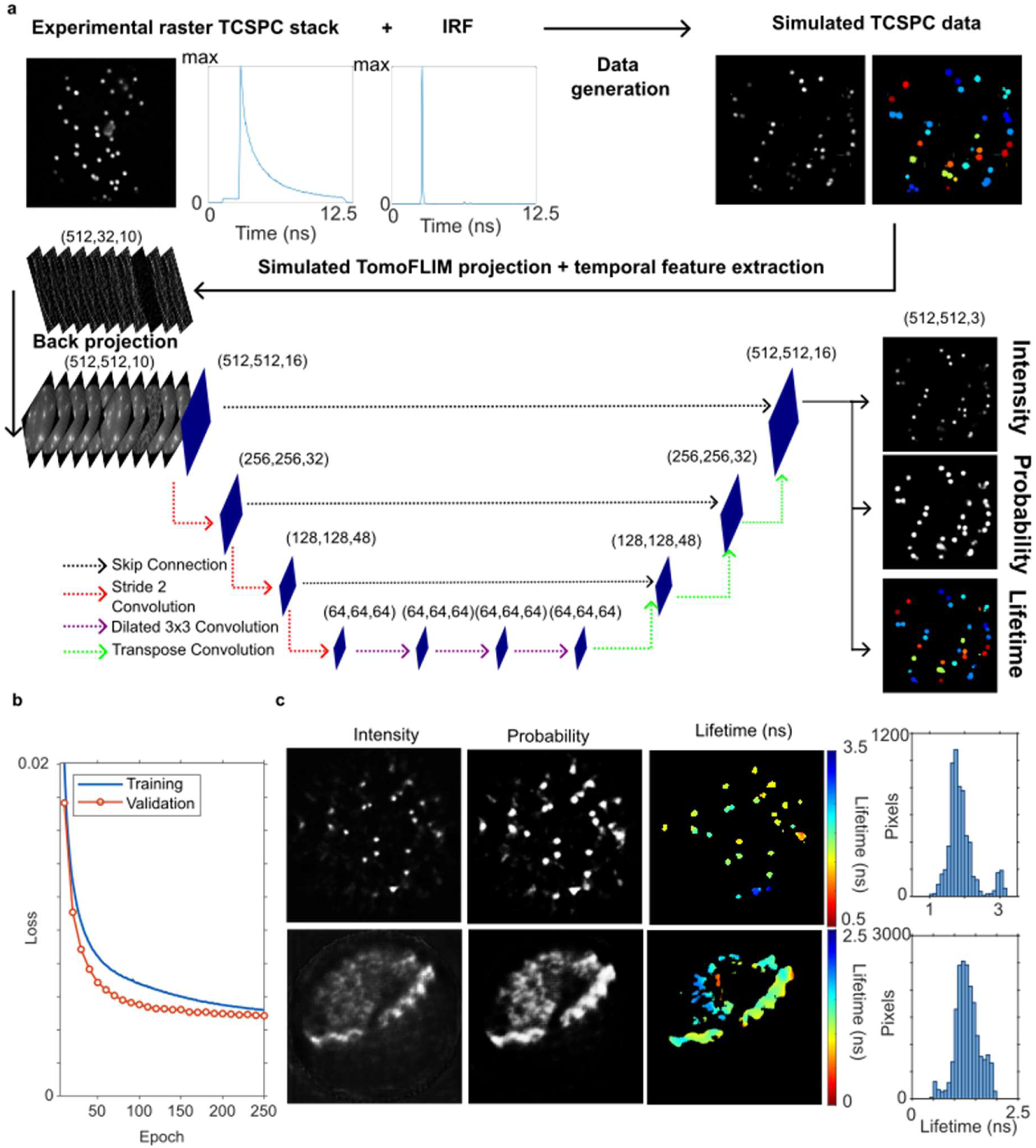
– Neural-network reconstruction of TomoFLIM data. **a** Physically informed TomoFLIM Net pipeline, including simulation-based training-data generation, temporal feature extraction, backprojection, encoder–decoder CNN reconstruction, and simultaneous intensity, fluorophore-probability, and continuous-lifetime outputs. **b** Epoch-wise training and validation loss over 250 epochs. **c** Representative experimental TomoFLIM reconstructions showing relative intensity, fluorophore probability, masked continuous lifetime, and the corresponding lifetime distributions.

Having established the quantitative performance of TomoFLIM Net on the test data, application of the trained network to experimentally acquired 32-angle TomoFLIM projections produced spatially resolved intensity and lifetime maps with clear lifetime contrast between fluorescent structures. **Figure 7c** shows TomoFLIM Net reconstructions of projections of both beads and Convallaria. The beads show clearly well-defined populations of each fluorescent lifetime species and excellent spatial similarity to the raster-scanned and CMM data shown in **figure 5**. Whilst the spatial likeness of the Convallaria has been well represented, the lifetime is clearly less reliable and bead-like artefacts are visible. This is likely because **t**he current network was trained predominantly on sparse, bead-like morphologies and therefore provides its strongest performance for punctate fluorescent samples. Nevertheless, these results demonstrate that physically informed neural reconstruction can provide a direct route from compressed TCSPC projection data to quantitative lifetime maps. Extension of the training set to include more diverse cellular and tissue-like structures should further improve applicability to complex biological specimens.

## Discussion and conclusion

The results demonstrate that TomoFLIM provides a clear route to accelerating multiphoton fluorescence lifetime imaging by reducing the redundant information acquired whilst retaining sufficient photon statistics for measurement of fluorescent lifetime. Conventional raster-scanned TCSPC-FLIM achieves excellent temporal resolution and lifetime accuracy, but results in acquisition of large quantities of redundant information. In contrast, parallelised SPAD-array approaches increase throughput by acquiring multiple spatial locations simultaneously, but require spatially resolved fluorescent signals and can introduce limitations associated with detector efficiency and broad wavelength dependent instrument response functions. Compressive single-pixel FLIM addresses the measurement bottleneck by encoding spatial information into a reduced number of measurements, although existing implementations have focused on wide-field imaging and do not provide the optical sectioning of multiphoton excitation. TomoFLIM occupies an intermediate regime between these approaches: by combining diffraction-limited multiphoton line excitation with single-pixel TCSPC detection, it retains the optical sectioning and high temporal resolution of conventional multiphoton FLIM while substantially reducing the number of spatial measurements required for image reconstruction. TomoFLIM could also potentially be extended to other fluorescent imaging modalities where acceleration of imaging frame rates is desirable. By frequency upconverting the laser (or changing to a picosecond pulsed diode) and moving the detector into the de-scanned path, the system could be adapted for confocal fluorescence lifetime imaging.

TomoFLIM can achieve compression ratios exceeding 90% while preserving spatial structure and lifetime contrast. A 32-angle projection (which represents a compression ratio of 93.75%) could reconstruct the intensity of Convallaria with PCCs of over 0.8 and sharper lifetime histograms than recorded in raster-scanned mode. Importantly, the dwell time was consistent between TomoFLIM and raster-scanned imaging. This meant that for a raster-scanned image acquired over 60 s, an equivalent TomoFLIM dataset was recorded with 3.75 seconds of live illumination. In addition, to allow the cylindrical lens to rotate to a new angle, the measurement was paused for approximately 0.1 s per angle (i.e. a 32-angle projection will require 3.2 seconds of dead time where the sample is not illuminated). In principle, by adopting alternative projection and beam shaping techniques, this dead time could be removed.

In addition to the CMM approach, the implementation of TomoFLIM Net demonstrates the potential for neural networks to further accelerate the reconstruction of compressed fluorescence lifetime data. A key advantage of this approach is that relative fluorescence intensity, fluorophore probability and continuous fluorescence lifetime can be recovered simultaneously from a single set of physically interpretable input features. This avoids the requirement for repeated iterative reconstruction and subsequent lifetime estimation, potentially allowing the computational reconstruction time to approach the acquisition time of TomoFLIM. The inclusion of a fluorophore-probability output is particularly useful for compressed lifetime imaging, as it provides an independent measure of where sufficient fluorescence signal is present and enables unreliable lifetime values in background regions to be rejected. Furthermore, because the network is trained using the experimentally measured instrument response function and illumination patterns, the reconstruction can incorporate characteristics of the physical imaging system directly into the learned mapping between the compressed measurement and reconstructed image. The strong reconstruction of the fluorescence lifetime calibration beads demonstrates the potential of this approach. The performance on more structurally complex samples should improve as the training dataset is expanded to include a broader range of biological morphologies, providing a route towards rapid reconstruction of TomoFLIM data for live-cell imaging.

A potential limitation of TomoFLIM is its sensitivity to pulse-pile-up in RSS-TCSPC measurements. In the present implementation, the dwell time is approximately 200 μs, corresponding to 16000 pulses at 80 MHz. Following the 5% rule^37^, this would permit an upper limit of 800 counts per pixel (in projection space) before pile-up related lifetime distortion becomes significant. Whilst the measurements presented in this article do not exceed the pile-up threshold, applications with higher count rates or laser powers would likely start to experience issues associated with pulse pile-up. One potential route to increasing the available count-rate capability would be the use of parallelised SPAD detection, in which the detected fluorescence is distributed across multiple independent detector elements. Such an implementation could also retain additional spatial information in the detected fluorescence, which could potentially be incorporated into the reconstruction process.

These characteristics make TomoFLIM particularly well suited to applications where rapid acquisition of quantitative lifetime information is required. In live-cell microscopy^38^ and in-animal imaging^17,39^, the reduced acquisition time could enable imaging of dynamic processes such as intracellular signalling, vesicular trafficking, and morphological changes while reducing temporal averaging and motion artefacts. The combination of multiphoton excitation and lifetime contrast may also be advantageous for Förster resonance energy transfer (FRET) measurements, where rapid acquisition could facilitate the study of transient molecular interactions. Furthermore, the optical sectioning provided by multiphoton excitation could enable faster lifetime imaging in thick or optically scattering specimens, including tissue and organoid models^40^, where conventional raster-scanned FLIM is limited by acquisition speed and sample-induced aberrations. More broadly, the ability to rapidly acquire spatially resolved lifetime information could extend quantitative FLIM to biological processes that currently occur on timescales faster than conventional acquisition methods can capture.

## Materials and Methods

### Optical System

The design of the optical system is shown in **Figure 1**. Light from a tunable Ti:Sapphire laser (Coherent, Chameleon Vision S) passes through a variable attenuation (VA) unit and through a beam expander formed of two achromatic doublet lenses L1 and L2 of focal lengths 30 mm and 250 mm, respectively (Thorlabs - AC254-030-B-ML, AC254-250-B-ML). The light passes through a beam-path selector formed of two flip mirrors (FM1 and FM2). When FM1 is active, the light is focussed by a 200 mm spherical lens L3 (Thorlabs - AC254-200-B-ML). When FM1 is out, the beam is focussed by a 200 mm achromatic doublet cylindrical lens (RCL, Thorlabs - ACY254-200-B) mounted in a rotary unit (Thorlabs - ELL14K). The light focussed by the cylindrical lens will pass to the microscope module when FM2 is in. RCL and L3 are positioned in conjugate planes and form a telescope with the 100 mm spherical achromatic lens L4 (Thorlabs - AC254-100-B-ML). The light is reflected by dual-axis galvanometric mirrors (Thorlabs – GCM102/M) and through an optical aperture. A removable, 50:50, non-polarizing beam splitter (BS, Thorlabs - BS014) is positioned after the optical aperture for visualisation of the excitation light on CMOS (Thorlabs-DCC1545). The light passes through the aperture and 100 mm scan lens L5 (Thorlabs - AC254-100-B-ML) into the microscope. The light is collimated by a 200 mm tube lens L6 (Thorlabs - AC254-200-B-ML) and transmitted through the dichroic (Semrock) and into the objective (Nikon, Plan Fluor, 40x/1.30, oil). The fluorescence is collected by the objective and reflected by the dichroic into the collection path. It is resized by a telescope formed of lenses L7 (Thorlabs - AC254-150-A-ML) and L8 (Thorlabs - AC254-100-A-ML) to match the size of the detector. A removable fluorescence filter was positioned before the hybrid detector (HPM-100-40), all data was collected with 920 nm excitation and a green emission filter in place.

The photon arrival times were measured on the TCSPC unit in RSS mode (Becker & Hickl - SPC 830), which was connected to the laser trigger output (80 MHz) with a pulse inverter attached directly to the SPC830. Both tomography and raster-scanned imaging were performed in FIFO imaging mode in the manufacturer’s software^5^. Acquisition was synchronised using a data acquisition device (National Instruments – NI-6363 DAQ) connected to the galvo-scanners and SPC830. The cylindrical lens was controlled directly from the computer by a USB connection to rotate the lens clockwise by angle *θ* relative to the y axis in the image plane. In raster-scanned imaging mode a linear voltage ramp was applied to the galvo-mirrors and in tomographic mode the line beam was projected along an angle orthogonal to *θ*, mathematically, this can be expressed as,

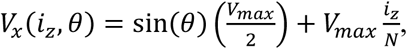

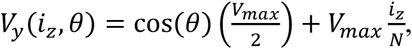

where *i_z_* is the number of acquisition points along the axis of projection, *V_max_* is the voltage value giving the full ramp between the spatial extrema of the field of view. In addition, a timing signal is also sent from the DAQ to the TCSPC module to synchronise the fluorescence measurement with the galvanometer mirrors. In tomographic mode, the x clock is synced with *i_z_* and the y clock with *θ*. The timing signal, the voltage ramp and rotation of the cylindrical lens is applied to the DAQ using customised MATLAB 2025 software developed in-house.

### Measuring lifetime with curve fitting

Raster-scanned scanning images were analysed using a curve fitting approach in the software package TRI2^34^. Cursors were estimated from the instrument response function, and baseline values were estimated with positivity constraints. A threshold was applied to the images, and the images were subject to a spatial kernel to ensure several thousand photons were in each fitted curve. Single exponential curves were fitted to recover the lifetime τ. The same approach was applied to the tomographic curve-fitted images shown in **Figure 3**.

### Lucy-Richardson reconstruction

The implementation of Lucy-Richardson reconstruction utilises the MATLAB open-source software package provided within^33^. This was used to perform reconstructions at either each time bin or of the first and zeroth moments of the CMM. To reconstruct an image, both the projection and a matrix containing the illumination patterns must be input to the software. For a *N* × *N* image *I* and equivalent projection *I_proj_* containing *N* ∗ *N_θ_* data points, the illumination patterns are also *N* × *N* matrices and are linearised and input to the illumination matrix *M,* which has dimension *N*^2^ × *N* ∗ *N_θ_*. This can be used to back-project the projection by matrix multiplying *M* and *I_proj_*.

### TomoFLIM Net

A convolutional neural network (TomoFLIM Net) was developed to reconstruct relative fluorescence intensity, fluorophore probability, and continuous fluorescence lifetime directly from time-resolved tomographic projection data. The network was trained using simulated (512 by 512) image scenes containing randomly positioned fluorescent objects, derived from the morphology of the raster-scanned image shown in **figure 4**. Connected fluorescent structures were extracted from this image to form a library of shapes, while their positions were randomised independently for every simulated scene. Objects were randomly rotated in increments of 90 degrees, reflected, and positioned subject to a non-overlap constraint and to the experimentally measured pattern-defined field of view. Each object was assigned an independently sampled mono-exponential lifetime, continuously distributed between 0.5 and 4.0 ns. The temporal response at each fluorescent pixel was generated by convolving the corresponding exponential decay with the experimentally measured instrument response function. The resulting time-resolved images were then forward-projected through the experimentally defined illumination matrix (32 angle illumination) and Poisson noise generation. A total of 1,250 simulated scenes were generated. A deterministic permutation with seed 1 assigned 1,000 scenes to training, 125 to validation and 125 to testing. Candidate networks were compared using validation lifetime mean absolute error.

For each acquisition, the measured projection data *I_i_*, were compressed into ten temporally informative measurements. These comprised the integrated signal, ∑*_i_I_i_*, normalised first temporal moment, 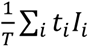 and eight temporally gated photon-count measurements, ∑*_i∈T_k__ I_i_* where *T_k_* denotes the *k*^th^ temporal gate. Each measurement was transformed to image space by backprojecting this dataset. The resulting ten (512 by 512) images formed the network input. To reduce sensitivity to variation in total photon counts between simulations and experiments, all feature channels were divided by the 99.5th percentile of the positive pixels in the \(M_0\) adjoint reconstruction.

The final network was trained using the Adam optimiser with a constant learning rate of 10^-4^, a mini-batch size of 16 and 250 epochs. Validation was calculated every 10 epochs, and candidate networks were compared using mean validation lifetime error. The encoder-decoder architecture used channel widths of 16, 32, 48 and 64, with learned stride-2 convolutions downsampling from 512 by 512 to 64 by 64. Two 64-channel bottleneck convolutions used dilation factors of 2 and 4. Transposed convolutions restored the image resolution, with additive skip connections between corresponding encoder and decoder scales. Layer normalisation followed encoder and bottleneck convolutions and decoder skip additions; hidden activations used Leaky-ReLU with a negative slope of 0.01, and dropout of 0.10 was applied at the bottleneck. The final (1 ×1) convolution produced three output logits. Sigmoid transformations were used to obtain relative intensity 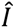, fluorophore probability 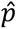 and continuous lifetime 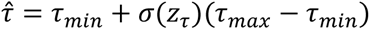. The network was then trained using a combined loss function:

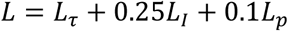

where *L_c_* and *L_I_* were mean-squared errors for lifetime and relative intensity, respectively, and *L_p_* was binary cross-entropy for the fluorophore mask. Lifetime loss was evaluated only at fluorescent ground-truth pixels. Convolutional weights were regularised with a coefficient of 0.01, and gradients were clipped to a global norm of 5. Training was performed in MATLAB 2024b using the Deep Learning Toolbox on an NVIDIA GeForce RTX 3060 Laptop GPU. The selected 250-epoch run required 4.31 hours; inference for one 32-angle experimental projection remained under 1 second.

For experimental TomoFLIM data, the measured projection cube was first arranged as detector position × time bin × projection angle. A detector- and angle-specific late-bin baseline was estimated from temporal bins 225–256 and subtracted before feature formation. The same temporal feature extraction, adjoint reconstruction, and per-acquisition normalisation were then applied before inference. Lifetime values were displayed only where the predicted fluorophore probability exceeded a predefined threshold. An empirical affine lifetime calibration, determined from tomographic reference measurements, was applied to the network output. This procedure enabled reconstruction of spatial intensity, fluorophore localisation confidence, and continuous lifetime maps from experimental time-resolved tomographic measurements without requiring arbitrary rescaling of photon counts.

## Data availability

The data that support the findings of this study are available from the corresponding author upon reasonable request.

## Author Contribution Statement

L.C., S.A.B, and S.P., conceived the project and designed the research. S.A.B, S.C., and S.P., supervised the project and acquired funding. L.C. built the instrument, developed the software for acquisition of TomoFLIM data and performed the data analysis and simulations of the experimental technique. L.C. and F.R. collected and curated all TomoFLIM and raster-scanned data. L.C., R.G.A., S.C., and S.P., contributed to design of the neural network. L.C. and S.P. wrote the manuscript. All authors reviewed and edited the manuscript.

## Ethics declarations

Conflict of interest - The authors declare no competing interests.

## Acknowledgements

This work was jointly funded by the Wellcome Trust Bioimaging Technology Development Awards and a UKRI Future Leaders Fellowship. Siân Culley is supported by the Royal Society University Research Fellowship. Fahmida Rahman is funded under an MRC Doctoral Training Programme.

