## Supplemental Text and Figures for "Multiphoton tomographic fluorescence lifetime imaging microscopy – TomoFLIM"

### Supplementary Information - Multiphoton tomographic fluorescence lifetime imaging microscopy – TomoFLIM

#### Simulating TomoFLIM

By generating simulated TCSPC data, it is possible to model the entire TomoFLIM process under different experimental conditions. A custom MATLAB script generated non-overlapping fluorescent bead distributions with defined fluorescence lifetime and photon-count characteristics. Beads were randomly positioned within the image and each bead was assigned a fluorescence lifetime randomly selected from a predefined set of lifetime species. A Gaussian spatial intensity profile was applied to each bead, with small random variations in peak brightness to introduce realistic inter-object intensity variability. For each pixel within a bead, the fluorescence photon arrival-time distribution was modelled as a normalised mono-exponential decay corresponding to the assigned ground-truth lifetime, with the total expected photon count determined by the local bead intensity. The data could then be projected to the resultant TomoFLIM data by multiplying the image at each time gate by the set of illumination patterns. This is the same illumination super matrix used in the Lucy-Richardson reconstruction. Poisson-distributed detector noise was subsequently added to the simulated fluorescence signal in both the image and the TomoFLIM projection. The resultant datasets thus had controllable sparsity, noise and distribution of lifetime counts. We were then able to compute the CMM of the raster images as well as the lifetime and intensity reconstructions from the simulated TomoFLIM data.

Figures S1-S4 show representative simulated TomoFLIM reconstructions for projections with 4, 8, 16 and 32 angles. The simulated TCSPC datasets had dimensions $128 \times 128 \times 128$ which corresponds to compression ratios of 96.88% ,93.75% ,87.5% and 75%. The bead intensity had a maximum of 4000 and on average there were 5 noise photons per pixel. The coverage ratio of the images varied from 1% to 5%. In total, 100 images were simulated and reconstructed at each compression ratio with 10 examples shown within this supplement. Due to the high signal to noise ratio, the computation of the CMM on the raster images matches extremely well with the simulated data (albeit with a slight bias at the longer lifetime). This effect is typical of CMM lifetime computation and can be compensated using calibration against a ground truth [1]. The relationship between sparsity and intensity/lifetime reconstruction is evident from both the PCC plot shown in the main text and observing the images shown herein. For the 4 angle projection, intensity and lifetime are only reconstructed faithfully for the sparsest of images we have investigated with broad, overlapping histograms. At lower sparsity, spatial features are lost and the histogram is an average of all lifetimes present in the simulated TCSPC data. By contrast at 16 and 32 angle projections, even the least sparse images exhibit strong visual similarity to the ground truth and CMM images shown above. The histograms also exhibit clearly defined populations of each lifetime species.

The simulation framework was also used to investigate the effect of increasing background on TomoFLIM reconstruction (Figure S5). For this analysis a 16 angle TomoFLIM acquisition was simulated while mean background contribution was increased from 1 to 100. For the raster images, this increases the overall noise level in the data from 16,384 to 1,638,400 whilst for TomoFLIM only 2048 to 204,800. Whilst the CMM computation on the raster data clearly deteriorates with increasing noise (evident by the broadened histograms), the TomoFLIM result is not visibly affected.


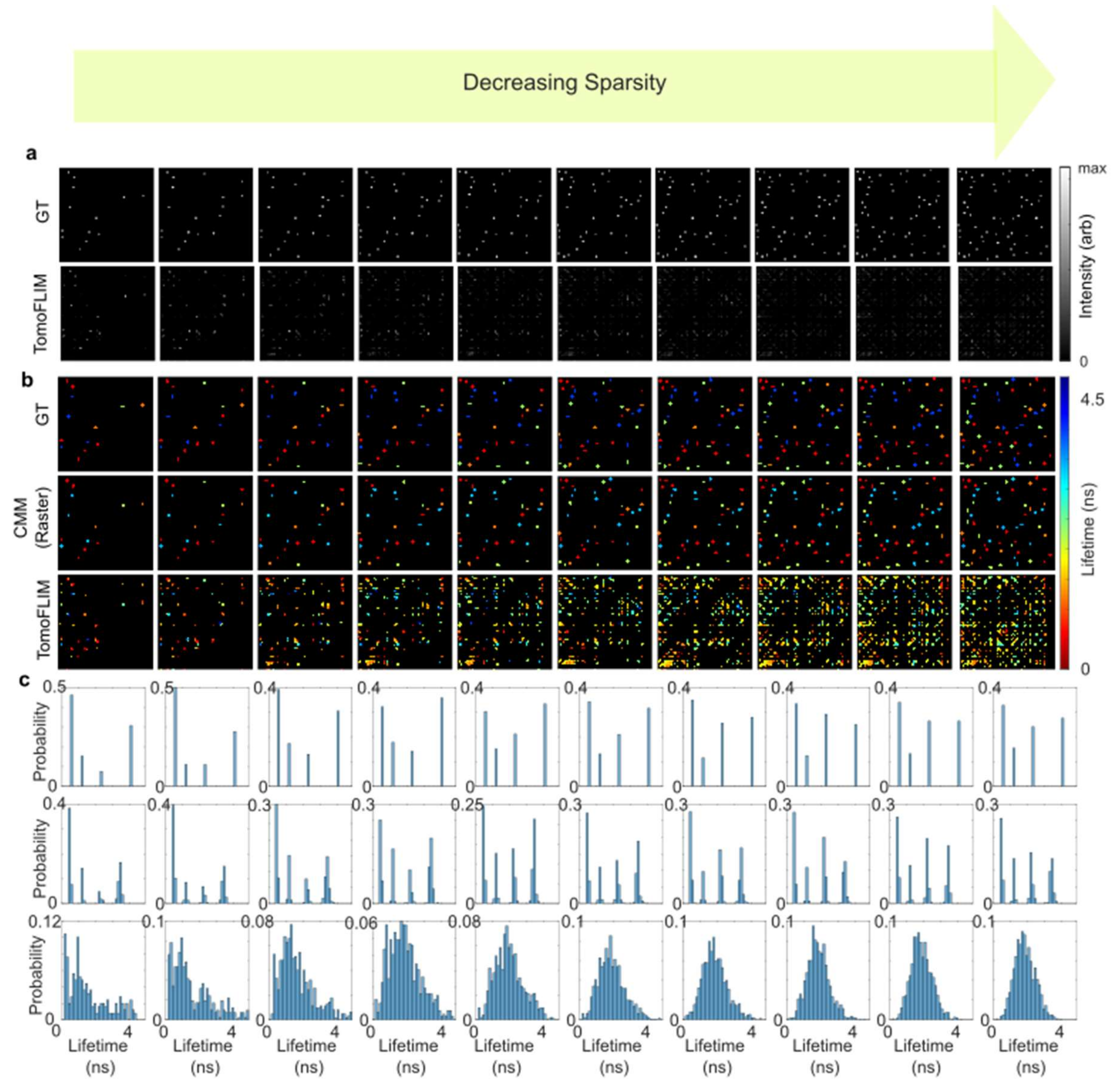


*Figure S1) – 4 Angle Projection **a** Intensity Ground Truth (top) intensity from TomoFLIM (bottom). **b** Lifetime ground truth (top), calculated CMM on raw TCSPC data (middle), CMM calculated on simulated TomoFLIM **c** Histograms of b.*

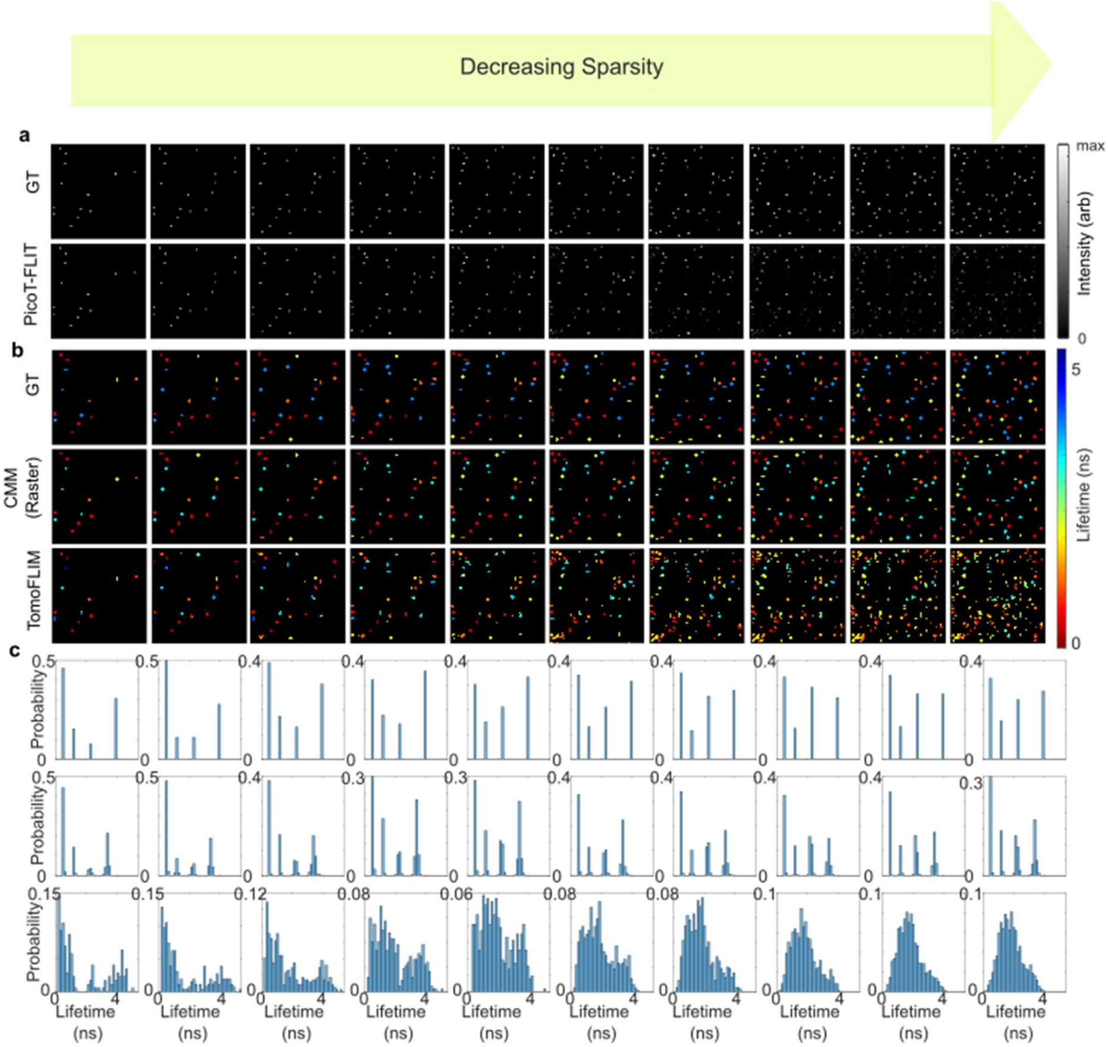


*Figure S2 - 8 Angle Projection A) Intensity Ground Truth (top) intensity from TomoFLIM (bottom), B) Lifetime ground truth (top), calculated CMM on raw TCSPC data (middle), CMM calculated on simulated TomoFLIM C) Histograms of B.*

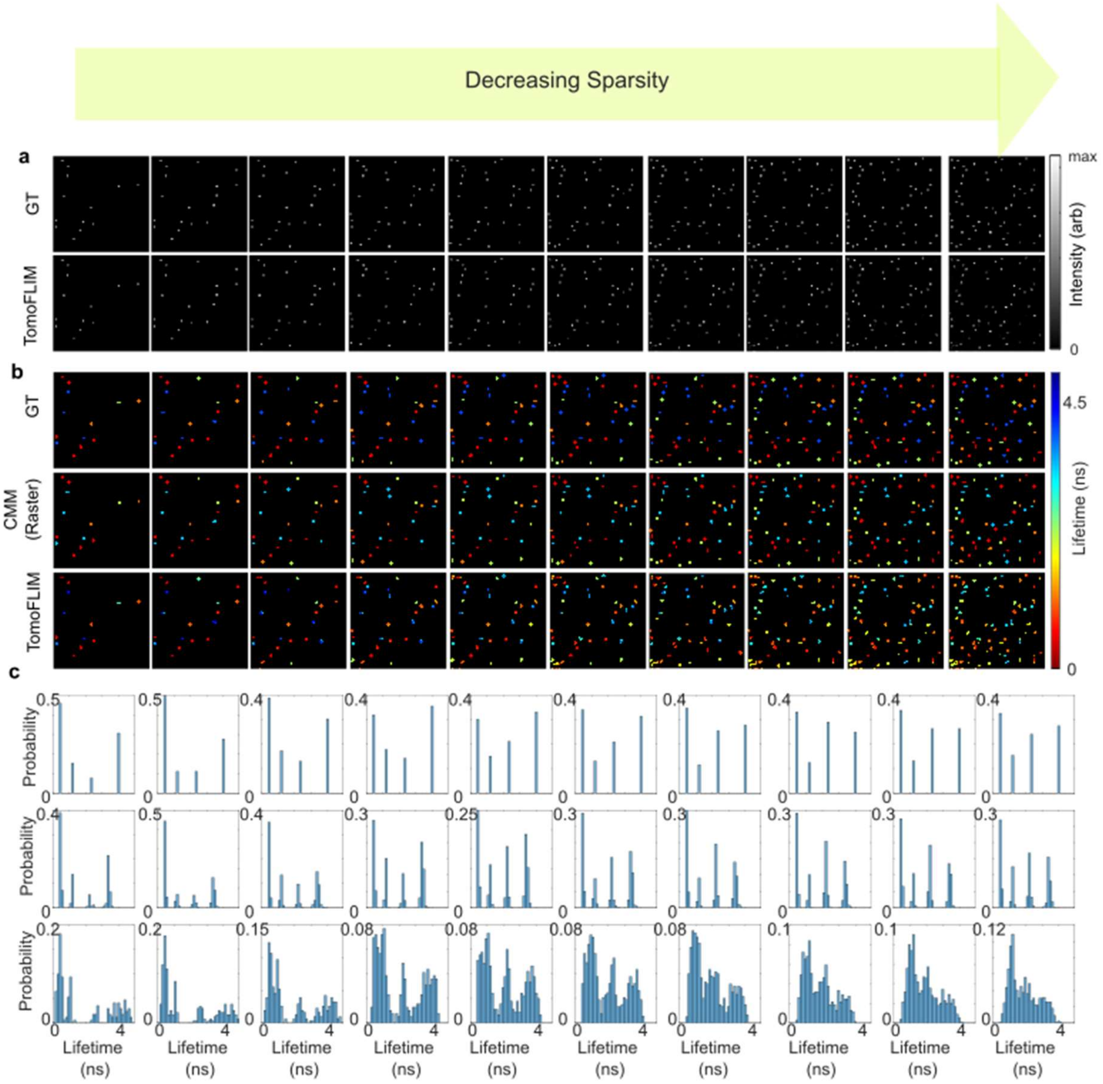


*Figure S3 - 16 Angle Projection A) Intensity Ground Truth (top) intensity from TomoFLIM (bottom), B) Lifetime ground truth (top), calculated CMM on raw TCSPC data (middle), CMM calculated on simulated TomoFLIM C) Histograms of B.*

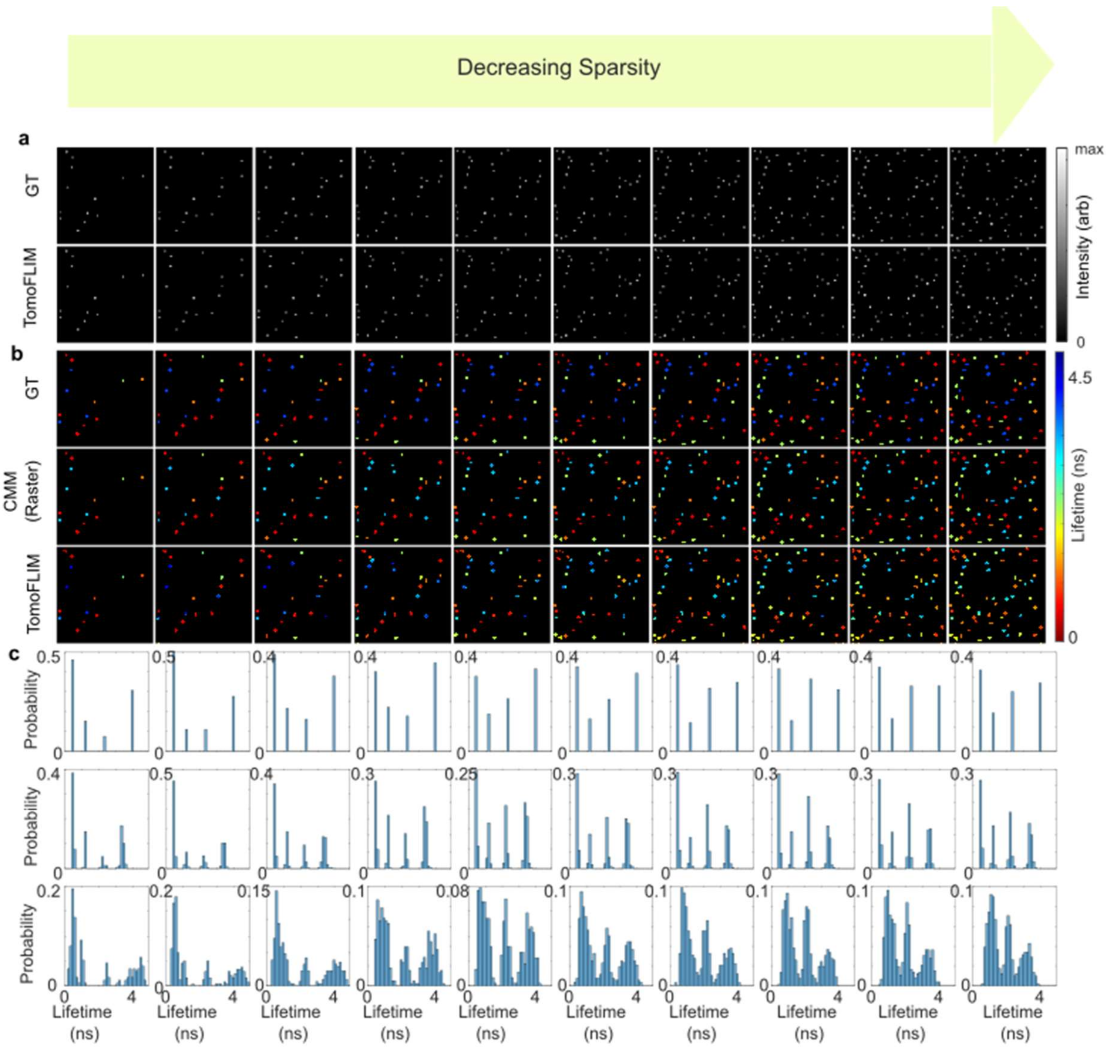


*Figure S4 - 32 Angle Projection A) Intensity Ground Truth (top) intensity from TomoFLIM (bottom), B) Lifetime ground truth (top), calculated CMM on raw TCSPC data (middle), CMM calculated on simulated TomoFLIM C) Histograms of B.*

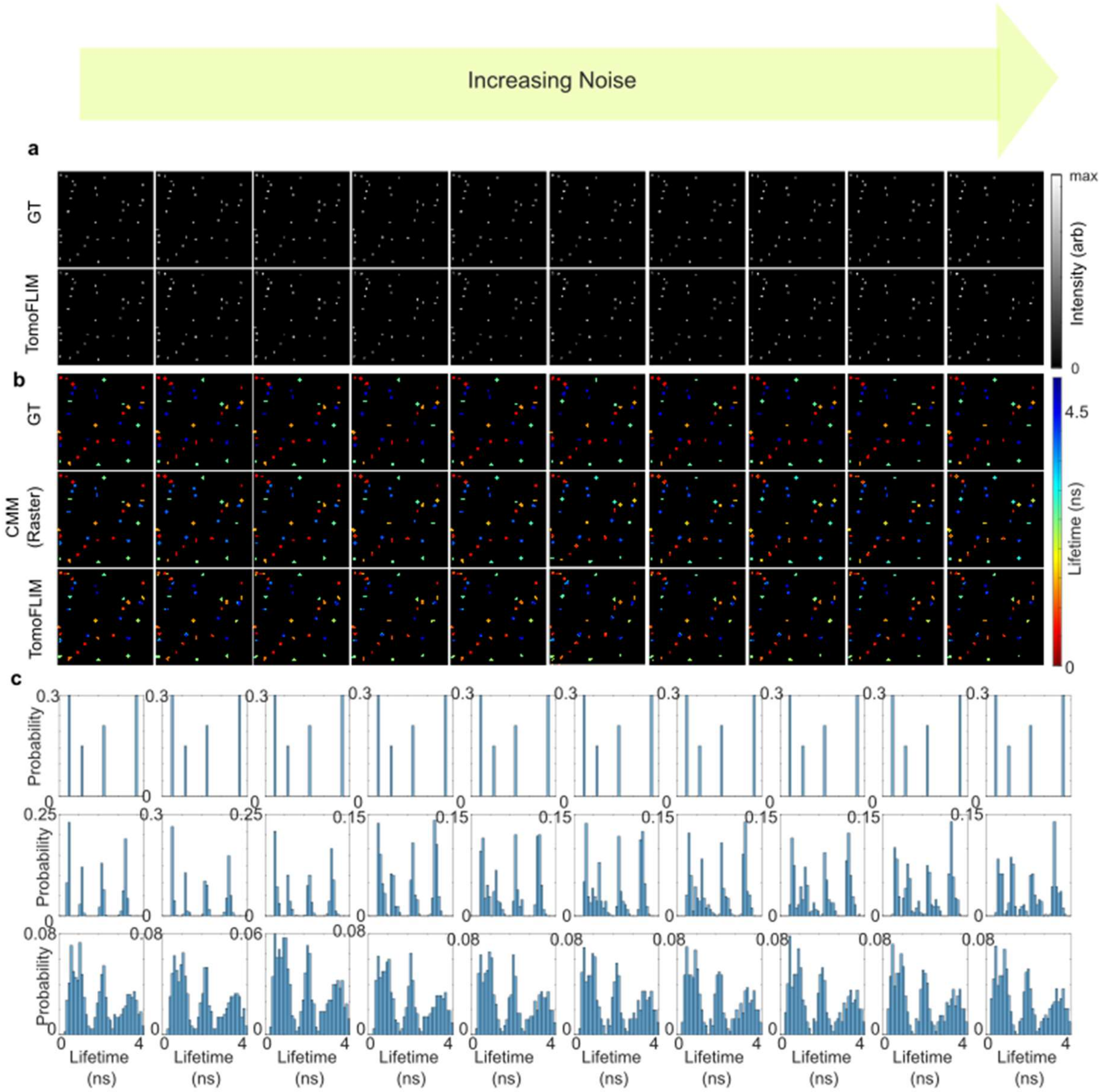


*Figure S5) 16 Angle Projection for data with varying noise A) Intensity Ground Truth (top) intensity from TomoFLIM (bottom), B) Lifetime ground truth (top), calculated CMM on raw TCSPC data (middle), CMM calculated on simulated TomoFLIM C) Histograms of B.*

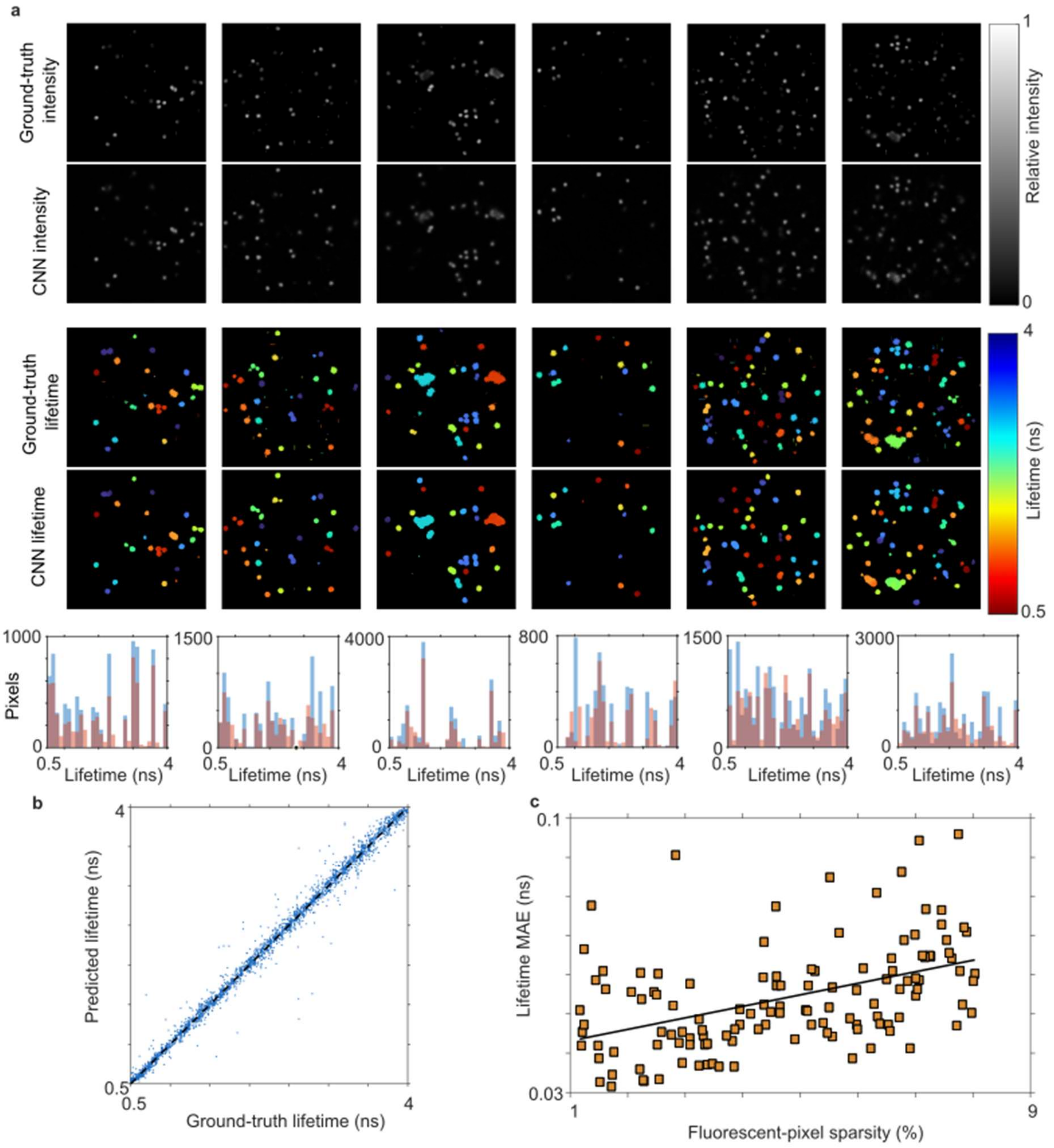


*Figure S6) Performance of TomoFLIM Net on simulated data.* ***a****, Reconstructions from six scenes selected at evenly spaced ranks of scene-level lifetime MAE. Rows show ground-truth and CNN-reconstructed relative intensity, ground-truth and CNN-reconstructed fluorescence lifetime, and the corresponding lifetime distributions. Intensity is shown on a common greyscale from 0 to 1. Lifetime maps and histograms use a 0.5–4.0 ns scale; background or masked lifetime pixels are black. Histogram colours denote ground truth (blue) and CNN prediction (orange).* ***b****, Agreement between ground-truth and CNN-predicted lifetime for 3,000 foreground pixels sampled across the internal synthetic benchmark. The dashed line indicates unity.* ***c****, Per-scene lifetime mean absolute error as a function of ground-truth fluorescent-pixel sparsity. The line denotes a least-squares linear fit.*
